# Globally distributed bacteriophage genomes reveal mechanisms of tripartite phage-bacteria-coral interactions

**DOI:** 10.1101/2024.03.11.584349

**Authors:** Bailey A. Wallace, Natascha S. Varona, Poppy J. Hesketh-Best, Cynthia Silveira

## Abstract

Reef-building corals depend on an intricate community of microorganisms for functioning and resilience. Bacteriophages are the most abundant and diverse members of these communities, yet very little is known about their functions in the holobiont due to methodological limitations that have prevented the recovery of high-quality viral genomes and bacterial host assignment from coral samples. Here, we introduce a size-fractionation approach which increased bacterial and viral recovery in coral metagenomes by 9-fold and 3-fold, respectively, and enabled the assembly and binning of bacterial and viral genomes at relatively low sequencing coverage. We combined these viral genomes with those derived from 677 publicly available metagenomes, viromes, and bacterial isolates from stony corals to build a Global Coral Virome Database of over 20,000 viral genomes and genome fragments spanning four viral realms. The tailed bacteriophage families *Kyanoviridae* and *Ackermannviridae* were the most abundant, replacing the since-abolished groups Podoviridae and Siphoviridae. Prophage and CRISPR spacer linkages between these viruses and 626 bacterial metagenome-assembled genomes and bacterial isolates showed that most viruses infected Alphaproteobacteria, the most abundant class, and less abundant taxa like Halanaerobiia and Bacteroidia. A host-phage-gene network identified keystone viruses with the genomic capacity to eavesdrop and modulate bacterial quorum sensing, interfere with sulfur cycling, and direct molecular interactions with eukaryotic cells through the release of extracellular effectors. This study reveals the basis of bacteriophage roles in modulating ecological interactions not only among bacterial community members but also directly affecting tripartite interactions with the coral host and its endosymbiotic algae.

## Introduction

Microorganisms of the coral holobiont are fundamental to the ecological success of reef-building corals (1). A complex network of symbiotic interactions connects all coral holobiont entities, including viruses, bacteria, archaea, fungi, and dinoflagellates, broadening the functional capacity of the coral host, and influencing the way the coral adapts and interacts with the reef environment (2,3). In addition to the well-described coral-Symbiodiniaceae endosymbiosis, prokaryotic communities (bacteria and archaea) play crucial roles in coral health (4). These assemblages remain flexible in response to a changing environment and can mitigate the effects of environmental stressors (5). While we have gained a significant understanding of many coral-bacteria interactions, little is known about the roles of viruses in the holobiont. Metagenomic studies have shown that viruses, particularly double-stranded DNA viruses infecting prokaryotes (herein, phages), are the most prevalent and genomically diverse entities within the coral holobiont (6–10). In other holobionts, phages have been shown to modulate bacterial community composition (11–13) and defend against pathogens by adhering to mucosal surfaces and regulating bacterial colonization through lysis (14). In corals, the pathogen *Vibrio coralliilyticus* can weaponize the lytic cycle of prophages in its competitors by triggering induction, gaining a competitive advantage over other symbiotic and potentially commensal bacteria in corals (15). Phages have also been implicated in coral disease. For example, bleached and white plague-diseased tissues harbor distinct phage communities (16) and T4-like phages have been associated with black band disease mats (17). At the reef scale, other studies show that phages play a crucial role in shaping coral ecological interactions, including competitive dynamics for benthic space (18) and biogeochemical cycling (19).

Despite these advances in our understanding of phages in the coral holobiont, the extent of their contribution to holobiont functional diversity remains unknown, mostly due to technical limitations in obtaining high-quality viral genomes from corals (20–22). Coral host and Symbiodinaceae DNA overwhelmingly dominate shotgun metagenomes, preventing appropriate coverage of viral genomes necessary for assembly. Moreover, viruses have high sequence diversity, lack hallmark genes, and display high levels of recombination, further complicating the assembly and characterization of viral metagenome-assembled genomes (vMAGs) (10,23,24). Combined with the limited availability of reference viral genomes, most studies to date have relied on the analysis of read data without assembly of viral genomes (10,25). These studies also characterized the viral community using a taxonomy framework that uses phage tail morphology (families Myoviridae, Podoviridae, and Siphoviridae) and the sequence identity of genes in these families to classify unknown viruses. This classification framework was abolished by the International Committee on Viral Taxonomy (ICTV) in 2022 in favor of a taxonomy system based on genomic composition, which better captures the evolutionary histories of viral groups (26). Therefore, an updated perspective on the diversity and functional genomics of coral-associated viruses in light of current approaches and taxonomic framework is due.

Here, we introduce a method for enriching viruses and bacteria in coral-associated metagenomes. We combine 33 metagenomes generated with this approach with publicly available datasets in a meta-analysis totaling 710 coral metagenomes, viromes, and bacterial isolates to reveal the genomic repertoire of coral holobiont viruses, focusing on phages. From this robust dataset, we identified 20,397 viral genomes or genome fragments from 31 coral species distributed across seven oceanographic regions. The phage community was dominated by the class *Caudoviricetes* (tailed phages, realm *Duplodnaviria)* but also included *Monodnaviria* and *Varidnaviria* phages. The ability to assemble viral and bacterial genomes enabled, for the first time to the best of our knowledge, matching phage host pairs in coral microbiomes via CRISPR matches. Phages most often infected hosts in the Alphaproteobacteria, Gammaproteobacteria, and Bacteriodia classes and encoded 109 unique metabolic genes and 64 unique virulence genes with the potential to affect host metabolism and symbioses. These results transform our knowledge of the coral virome by identifying specific bacteria-phage-gene links involved in quorum sensing, sulfur cycling, and direct interactions with eukaryotes within the holobiont.

## Methods

### Development of a viral and bacterial enrichment method

Fragments of the coral *Orbicella faveolata* (N = 28) were collected via SCUBA in July 2021 along the southwestern coast of Curaçao in the Caribbean (Table S1). Specimens of approximately 1cm^3^ (containing mucus, tissue, and skeleton) were collected using a chisel and hammer and placed in Ziplock polyethylene bags. The specimens were briefly placed on ice during transfer to the laboratory at the CARMABI research station, where they were flash-frozen and stored at −80°C until later processing. The samples were subjected to two DNA extraction protocols for comparison: 19 samples were processed using the Viral and Bacterial Enrichment (VBE) method developed here, and 6 underwent bulk extraction and sequencing and were treated as controls. For both extraction methods, coral fragments were thawed, crushed to a fine gravel texture using a sterile mortar and pestle, and suspended in 150 µL of sterile artificial seawater. Control coral homogenates were extracted using a DNeasy PowerSoil Kit (QIAGEN, Germantown, MD, United States), modified with the addition of 20 µL of Proteinase K incubated at 56°C for 10 minutes prior to the kit’s lysis steps. For VBE, the coral homogenate was placed into a tube containing ∼ 0.2 g of 425-600 µm sterile glass beads (Sigma-Aldrich, St. Louis, MO, United States) and vortexed at speed 3 (∼600rpm) for 5 minutes (VWR Analog Vortex Mixer; VWR, Radnor, PA, United States). The supernatant was transferred to a clean tube, brought up to 1 mL with sterile ASW, and treated with DNAse I (20 U/mL final concentration) in DNAse I Buffer (Invitrogen, Waltham, MA, United States) at 25°C for 2 hours. DNAse activity was stopped with EDTA (0.005 M final concentration; Genesee Scientific, San Diego, CA, United States) and the sample was filtered through an 8.0 µm membrane (Cytiva, Marlborough, MA, United States). The flowthrough was collected, transferred to a 100 KDa Amicon Centrifugal Unit (Sigma-Aldrich), and centrifuged at 3,200 g for 30 minutes. After concentration, each side of the Amicon filter was rinsed and incubated at 56°C for 1 hour (per side) in 200 µL Buffer T1 and 20 µL Proteinase K from the NucleoSpin^l1J^ Tissue Kit (MACHEREY-NAGEL Inc., Allentown, PA, United States) and DNA extraction proceeded from step 3 of the kit. DNA was eluted in 100 µL of PCR-grade water and quantified with a Qubit 2.0 Fluorometer (Invitrogen). A step-by-step description of the VBE protocol is publicly available on protocols.io (27). Library preparation and whole genome sequencing were conducted by Azenta Life Sciences (South Plainfield, NJ, United States). Metagenomic libraries were generated with the NEBNext Ultra DNA Library Preparation kit following the manufacturer’s instructions (New England Biolabs, Ipswich, MA, United States) and paired-end sequenced (2 x 150 bp) on an Illumina HiSeq 4000 platform (Illumina, San Diego, CA, United States).

### Quantification of viral and bacterial enrichment in coral metagenomes

Raw metagenomic reads were adapter-trimmed, quality-filtered (trimq=30, maq=30), and entropy-filtered (entropy=0.90) using BBDuk (28), generating 594,487,714 quality-controlled reads. To identify coral host and Symbiodinaceae reads, the quality-controlled (QC) reads were mapped to the genome of the coral host, *O. faveolata* (GCA_001896105.1), and Symbiodinaceae (Clade A) (GCA_003297005.1) using Bowtie2 (--mp 4 and -X 1000) (29). Mapped reads were removed using SAMtools v1.18 (30) and quantified with SeqKit (31), yielding 336,288,370 coral and Symbiodinaceae filtered reads (Table S1). The QC reads and coral and symbiont filtered reads from each sample were assembled separately using metaSPAdes v3.15.5 with default parameters (32). Bacterial reads were identified with Kaiju v1.9.0 in each group using the Progenomes database (33) and normalized by either the number of QC reads or coral and symbiont filtered reads in each sample to obtain the percentage of bacterial reads in each metagenome. Bacterial MAGs were generated by integrating the single-sample coverage binning outputs from MaxBin2 v2.2.7 (34), MetaBat2 v2.15 (35), and CONCOCT v1.1.0 (36). The resulting bins were consolidated and improved with the metaWRAP v1.2.1 (37) bin refinement module. The quality of refined bins was assessed with CheckM2 v1.0.2 (38) to select bins with ≥50% completion and ≤10% contamination. VIBRANT v1.2.1 identified putative viral genome fragments among contigs (39). Viral contigs were pooled by DNA extraction method to create two databases, “VBE” and “Control”. The number of reads mapped to their respective databases at 80% identity with SMALT v0.7.6 (40) was normalized by the total number of quality and filtered reads and by the length of each contig (41) to obtain the viral fractional abundance (see Fig. S1 for bioinformatic pipeline). Comparisons between VBE and controls were based on Student’s t-test between the two groups, apart from Figure 1E, which displays a simple count of bMAGs generated from each group. Taxonomic assignment of phages was performed with Phage Taxonomy Tool (PTT) (42), with the reference taxonomy edited to reflect updated ICTV classifications (43).

**Figure 1.**
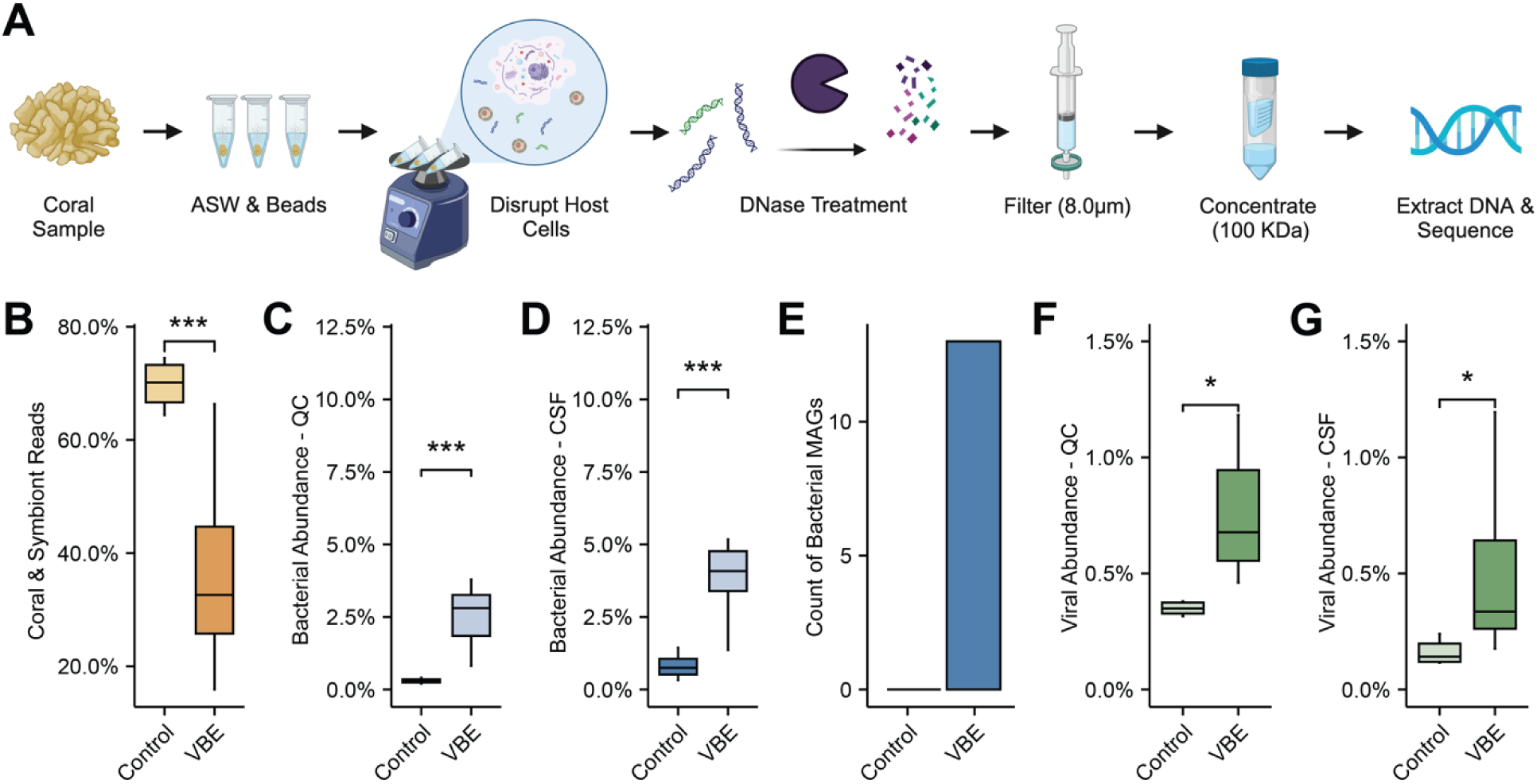
Enrichment of viruses and bacteria from coral samples. **(A)** Simplified workflow depicting the VBE method for enrichment of viruses and bacteria in coral metagenomes. **(B-G)** Recovery of coral and symbiont (orange), bacteria (blue), and viruses (green) in VBE and control metagenomes. Error bars represent **(B)** Percent of coral and symbiont reads, **(C)** Bacterial abundance within coral and symbiont-filtered (CSF) reads, **(D)** bacterial abundance within quality-controlled (QC) reads, **(E)** Abundance of bacterial MAGs, **(F)** Viral fractional abundance in CSF reads and **(G)** in QC reads. Stats represent p-values from statistical tests: Students t-test in panels B, C, D, F and G (* p < 0.05, ** p < 0.01, *** p < 0.001).

### Virus identification in global coral metagenome datasets

We assembled the Global Coral Virus Database (GCVDB) by combining viruses identified in the 28 Curaçao samples described above, 5 additional *O. faveolata* samples from Miami, FL (processed with VBE), and publicly available metagenomes (N = 355), viromes (N = 12), and genomes of bacteria (N = 310) isolated from Scleractinian corals (Tables S2 and S3 list the metadata and accession numbers for each dataset). Search terms and combinations of search terms including but not limited to “coral”, “metagenome”, “virome”, and “bacterial isolates” were used to identify publications and their associated datasets through Google Scholar, the National Center for Biotechnology Information (NCBI), the European Nucleotide Archive (ENA), and the JGI IMG/M Database (44). The inclusion of metagenomic data was restricted to Illumina sequences and excluded early 454 and Sanger sequencing studies.

The quality of the 310 bacterial genomes was assessed using CheckM v.2.0.12 (45). VIBRANT identified 637 putative viral genomes in these bacterial genomes. After dereplication with Virathon (46), 178 representative viruses were added to the GCVDB. Metagenomes and viromes contained 36,006,392,282 raw metagenomic reads that were quality-controlled and assembled with the same parameters described above. No coral or symbiont read filtering was performed on this larger dataset, as not all coral species had available reference genomes. Contigs longer than 1,000 bp were screened by VIBRANT and the identified viral genome fragments were grouped by project, location, and host coral species, and co-binned into viral metagenome-assembled genomes (vMAGs) with vRhyme v1.1.0 using the “longest” method which dereplicates scaffolds, keeping the longest representative sequence (47). Contigs within bins were subjected to further dereplication using Virathon and N-linked (1,000 Ns) within bins. N-linked vMAGs were dereplicated with Virathon, and their quality was assessed with CheckV v1.0.1 (48). This process generated 2,121 vMAGs and 18,098 viral contigs that were added to the GCVDB. Combined with the 178 viral contigs derived from bacterial isolates, a total of 20,397 unique putative viral genomes from coral holobionts compose the GCVDB (Table S4).

### Diversity of coral-associated viruses

Viral contigs annotated as “complete circular, “high-quality draft”, and “medium-quality draft” by VIBRANT (N = 317) were combined with vMAGs annotated as “high-quality” and “medium-quality” by CheckV (N = 528) for further analyses. These genomes were compared to the NCBI Viral RefSeq v1.1 database (accessed: 2023-07-13), which was first dereplicated using MIUViG-recommended parameters (95% average nucleotide identity, 85% AF) (49). The comparison was performed using GL-UVAB v0.6.pl, which calculates an all-versus-all Dice distance matrix based on the number of shared protein-encoding genes and their identity level (50). The distance matrix was used to build a virus proteomic neighbor-joining tree containing 845 GCVDB viruses and 257 NCBI viruses. The tree was visualized and annotated using Interactive Tree of Life (51). Taxonomic assignment of GCVDB viruses was performed by Phage Taxonomy Tool (PTT), which exclusively identifies prokaryotic viruses through protein similarity (42). The ‘*PTT_virus_taxonomy.tsv’* reference data sheet was edited to reflect updated ICTV classifications (43). Adonis and beta dispersion analysis was performed with the R package Vegan (52).

### Bacteriophage host identification

Bacterial MAGs were generated from all metagenomic samples using the methods described above. Briefly, single-sample coverage binning outputs from MaxBin2 v2.2.7 (34), MetaBat2 v2.15 (35), and CONCOCT v1.1.0 (36) were consolidated and improved with metaWRAP v1.2.1 (37), resulting in 910 bins. CheckM2 v1.0.2 (38) was used to select bins with ≥50% completion and ≤10% contamination. Bins with low confidence predictions were removed (N = 6), resulting in 316 bMAGs taxonomically classified with GTDB-Tk v2.3.2 (53) (Table S5). To predict virus-host pairs, we employed a combination of provirus detection and CRISPR spacer matching (Table S6). A database of CRISPR spacers from the 316 bMAGs and 310 bacterial isolates was generated with minCED v0.4.3 (54), a tool derived from CRISPR Recognition Tools (CRT) v1.2 (55). The resulting CRISPR spacer database was subsequently used to identify sequence homology matches with GCVDB using BLASTn. 63 high-confidence pairs were generated using thresholds of ≤ 2 mismatches/gaps, 100% coverage to the spacer, and a sequence length of ≥ 20 nucleotides. Provirus detection was accomplished by mapping the GCVDB viruses to bMAGs with Minimap2 v2.24-r1122 (56). Only the bMAG contigs that contained matches of 100% identity (no gaps) along the entire mapping length (N = 448) were selected to be assessed with CheckV, which identified 59 proviruses with host flanking regions. Viral contigs identified in bacterial isolate genomes by VIBRANT were categorized as proviruses (N = 177).

### Genetic repertoire encoded by phages

The genomes of the GCVDB viruses from metagenomic samples were compared to viruses identified in coral reef seawater metagenomes from the island of Curaçao (57). These water metagenomes are paired with a subset of the coral metagenomes described above (Table S1). KEGG gene annotations were used to identify potential metabolic genes (Table S7). We also searched for genes classified as virulence factors (VFs) encoded by viruses using a BLASTp comparison with a curated database of experimentally tested virulence genes (58) with an e-value cut-off of ≤0.00001 and ≥40% identity across ≥20 amino acids (Table S8). For multiple quality hits on overlapping regions, the best hit was selected based on the e-value. Only metabolic genes identified in genomes with taxonomic annotations by PTT were included in the analysis of phage-bacteria interactions. Simpson’s Index of Diversity and Pielou’s Evenness were used to compare the diversity and evenness of these groups.

### Tripartite network

A tripartite network of phage-host linkages and phage-gene linkages was constructed using Cytoscape v3.9.1 (59). To profile keystone constituents in this network, we treated shared metabolic and virulence genes as linkages between viruses and used the node degree (k; number of interactions), closeness centrality (cc; distance to all other nodes), and clustering coefficient (clust; connectivity of nearest neighbors) calculated by Cytoscape’s NetworkAnalyzer feature. We ranked each species considering all three properties and calculated an average rank to identify keystone viruses (60). Phage genomes with host linkages and genes of interest identified in this network were plotted with the R package genoPlotR (61).

## Results

### Virus and Bacteria Enrichment reduces coral and symbiont DNA in metagenomes

The Viral and Bacterial Enrichment (VBE; Fig. 1A) metagenomes had 36.39 ± 0.10% (mean, SE) coral or symbiont reads, compared to 69.83±0.13% (mean, SE) in the controls, a 33.44% (1.92x) reduction in the VBE metagenomes (t-test, t(23) = -5.58, p = 1.124e-05) (Fig. 1B). Among quality-controlled reads, 2.72±0.33% (mean, SE) of VBE reads were bacterial, compared to only 0.30±0.03% (mean, SE; here and hereafter) in controls, a 2.42% (9.01x) enrichment in VBE (t-test, t(23) = 4.12, p = 4.15e-04) (Fig. 1C). For coral and symbiont filtered reads, VBE increased bacterial recovery by 5.02x (4.06±0.09% and 0.81±0.06% for VBE and control, respectively; t-test, t(23) = 4.29, P = 2.73e-04; Fig. 1D). Remarkably, high quality bacterial metagenome-assembled genomes (bMAGs) could only be generated from VBE metagenomes, with 13 total bMAGs with ≥ 50% completion and ≤ 10% contamination identified (Fig. 1E). VBE generated 543 putative viral contigs, compared to 76 from controls, which represented 0.74±0.05% and 0.40±0.05% of QC reads, respectively, a 1.87x increase for VBE (t-test, t(23) = 3.45, P = 2.17e-03) (Fig. 1F). Among coral and symbiont-filtered reads, 0.46±0.06% were viral in VBE and 0.16 ± 0.02% in controls, a 2.83x increase in viral recovery by VBE (t-test, t(23) = 2.6878, P = 1.31e-02; Fig. 1G)

### An updated taxonomic profile of coral holobiont phages

The 710 publicly available metagenomes, viromes, and bacterial isolates from 31 coral species and seven ocean regions (Fig. 2A) analyzed here yielded 20,397 viral genomes or genome fragments, including viral contigs (vContigs) and viral metagenome-assembled genomes (vMAGs). These viral genomes were combined into the Global Coral Virus Database (GCVDB), and hereafter, we refer to these genomes and genome fragments as viruses for simplicity. A total of 846 viral genomes were classified as either medium quality, high quality, or complete circular genomes (hereafter, high and medium quality viruses). The proteomic tree in Figure 2B shows the relationships between high and medium-quality genomes in the GCVDB (N = 846) and viruses in the ICTV database (N = 99). GCVDB viruses span four viral realms according to the current ICVT classification: *Duplodnaviria* and *Varidnaviria* double-stranded DNA viruses, *Monodnaviria* single-stranded DNA viruses, and *Riboviria* RNA viruses similar to retroviruses with a DNA phase. Several families of ICTV viruses, including *Adintoviridae*, *Polydnaviriformidae*, *Caulimoviridae*, *Metaviridae*, and *Retroviridae*, which infect eukaryotes, as well as viruses obtained using the VBE method had representatives throughout this tree. Bacteriophages, specifically, belonged to three realms and 25 families spanning both dsDNA and ssDNA viruses (Table S9).

**Figure 2.**
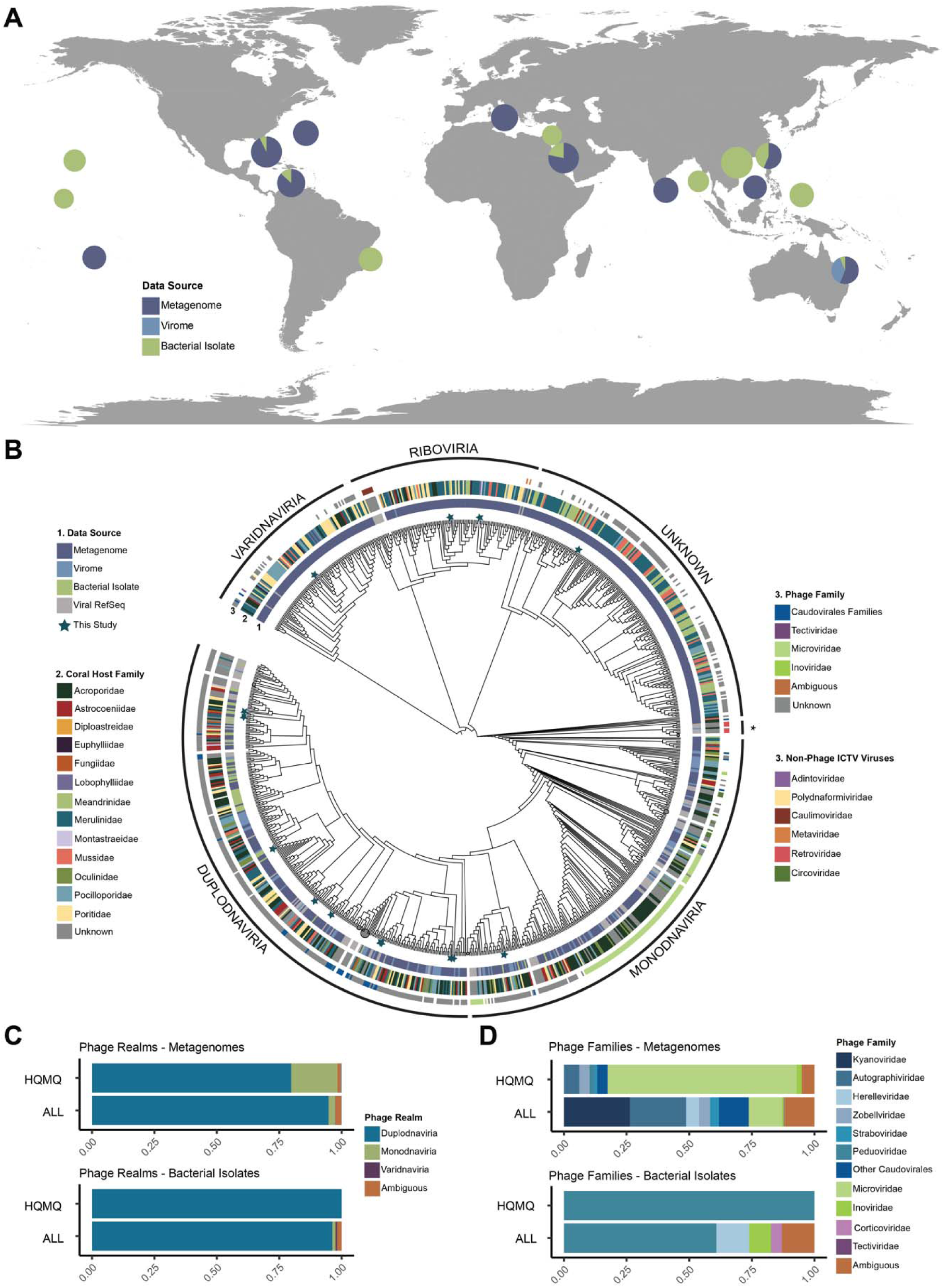
The coral virome diversity. **(A)** Distribution of sample locations and data types across ocean regions. Samples were grouped by oceanographic region for each pie chart. The radius of each pie chart represents the number of samples with log(_10_)+4 transformation. Colors indicate the type of dataset: metagenome, metavirome, or complete/draft bacterial genomes. **(B)** Proteomic tree of genomes and genome fragments in the Global Coral Virome database and their closest relatives from the NCBI Viral RefSeq. The innermost color ring (1) describes the data source (metagenome, virome, bacterial isolate), the second color ring (2) depicts the family-level taxonomy of the coral host where the virus was identified, and the outermost color ring (3) describes the family-level taxonomy of the viral genome (PTT taxonomy for phages and ICTV classification for RefSeq viruses). Viral realms were defined in the tree based on the position of RefSeq viruses. The branch labeled with an asterisk (*) contains Riboviria viruses. Branch lengths were omitted to better display the tree topology. Branches with three or more RefSeq members were collapsed (grey circles, proportionally sized to the number of collapsed nodes). Teal stars indicate 14 viruses obtained by the VBE method. **(C and D)** Taxonomic classification of phages in the complete GCVDB (ALL) and in the subset of this database containing viral genomes of high or medium quality (HQMQ) at the realm (**C)** family **(D)** taxonomic levels.

The majority (68.95%) of viruses across the GCVDB were not taxonomically classified due to the low similarity with reference viruses. Among the high and medium quality viruses in metagenomes and bacterial isolate genomes, most were identified as phages (58.85% and 97.96%, respectively). Of the phages with taxonomic annotations, *Duplodnaviria* was the dominant realm regardless of quality (Fig. 2C). In metagenomes, this was followed by *Monodnaviria*, which represented 2.67% of the classified phages in the full GCVDB and 16.20% of the high and medium quality phages. In the genomes of bacterial isolates, *Monodnaviria* were absent among high and medium quality but comprised 1.24% when including low-quality. 46.55% of *Duplodnaviria* viruses in the high and medium quality metagenomes did not have family classification under current taxonomy, resulting in an overrepresentation of ssDNA *Monodnaviria* families in Fig. 2D. While these *Monodnaviria* families, *Microviridae* and *Inoviridae*, were the most abundant families identified among the high and medium quality metagenomes, dsDNA tailed phages belonging to the class *Caudoviricetes* were dominant in all other groups. Several dsDNA families annotated as *Duplodnaviria* were more closely related to viruses from other realms in the proteomic tree. For example, viruses classified as *Straboviridae* and *Ackermannviridae* fell within the *Varidnaviria,* while *Peduoviridae* and other *Ackermannviridae* were grouped within the realm *Monodnaviria*.

We estimated the effects of coral host and ocean region in shaping virome composition by using the Jaccard beta diversity index. Viromes were significantly different across coral host families (*adonis*, R^2^ = 0.43, P = 0.001) and ocean regions (*adonis*, R^2^ = 0.36, P = 0.001). However, the project and subproject that generated the data explained the most variation in Jaccard’s beta diversity (60.78% and 71.99%, respectively). These variables are not independent of ocean region, as any given project was from a single ocean region, and each subproject includes both a single ocean region and a single coral species. The data displayed different levels of dispersion by ocean region (*betadisper*, F =4.36, P = 2.86e-04), coral host family (*betadisper*, F =15.71, P = < 2.2e-16), project (*betadisper*, F =18.32, P = < 2.2e-16), and subproject (*betadisper*, F = 7.2374, P = < 2.2e-16), indicating that the Adonis PERMANOVA results are affected by non-homogeneous dispersion of the data.

### Metabolic and virulence genes differ between coral and seawater viruses

The 3,912 seawater and 6,173 coral viruses classified here as phages encoded 181 unique genes with KEGG ortholog annotations (11.34% of coral and 27.30% of seawater phages). Seawater viruses encoded more unique metabolic genes (N = 72) than corals (N = 51), despite a >22-fold increase in sampling effort (18 seawater metagenomes were used here for comparison versus 400 coral metagenomes (Fig. 3A). Among the 58 metabolic genes shared between coral and seawater viruses, the most common were involved in amino acid metabolism, energy metabolism (Photosynthesis), and Sulfur relay (Fig. 3B). Only five of the ten most common metabolic genes in coral phages were also among the ten most common in seawater phages. Metabolic gene α-diversity, as expressed by Simpson’s Index of diversity, was higher in seawater (D = 0.95) than in corals (D = 0.61). Additionally, metabolic genes were more evenly distributed in seawater than in coral phages (E_Pielou_ = 0.72 in seawater and E_Pielou_ = 0.41 in corals), where a single gene, DNA cytosine-5 methyltransferase 1 (*DNMT1, dcm*), accounted for 60.35% of all metabolic genes identified. *DNMT1, dcm* was also the most abundant gene in seawater, yet only accounted for 13.25% of the metabolic genes. Together, the four most abundant metabolic genes in coral viruses, *DNMT1, dcm*, DNA cytosine-5 methyltransferase 3A (*DNMT3a*; 14.32%), S-(hydroxymethyl)glutathione dehydrogenase/alcohol dehydrogenase (*frmA*, *ADH5*, *adhC*; 3.29%), and nicotinamide phosphoribosyltransferase (*NAMPT*; 1.57%) accounted for 80% of the metabolic genes encoded by viruses in corals, while in seawater, 21 genes constitute that same 80% threshold. These common metabolic genes in coral phages are involved in sulfur-containing amino acid metabolism (*DNMT1, dcm; DNMT3a*), energy metabolism (methane metabolism; *frmA, ADH5, adhC*), and metabolism of cofactors and vitamins (Nicotinate and nicotinamide metabolism; *NAMPT*).

**Figure 3.**
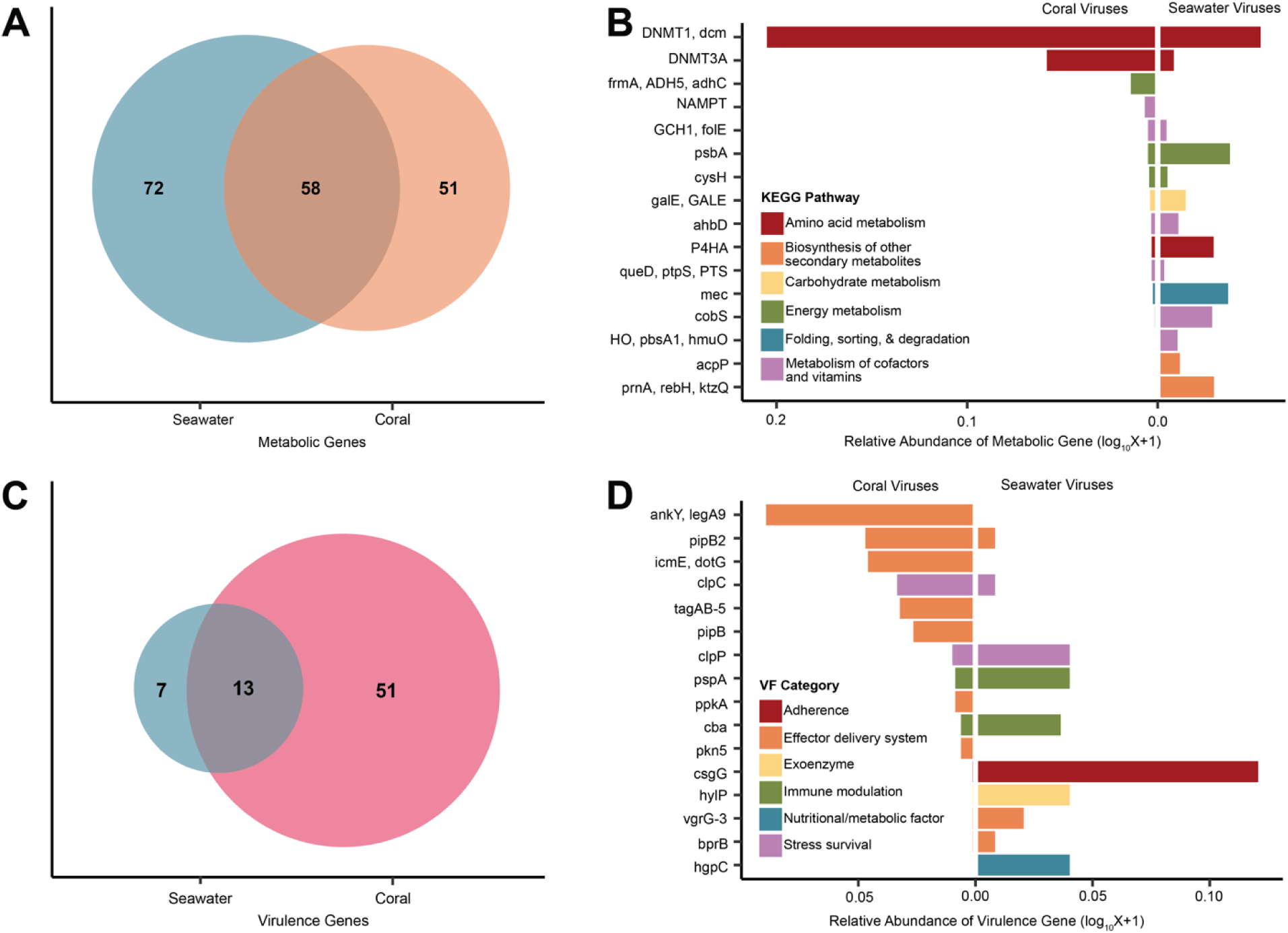
Genomic repertoire of coral-associated bacteriophages. **(A)** Metabolic genes (defined based on KEGG Orthologs) identified in coral and seawater bacteriophages. An overlap indicates those identified in both sample types. **(B)** Frequencies of the most common metabolic genes, calculated as the sum of viruses encoding each gene within each sample type and sorted by abundance. The colors indicate the KEGG pathways to which those genes belong. **(C)** Virulence factors identified in coral and seawater bacteriophages and **(D)** frequencies of the ten most common VFs, calculated as the sum of the phages containing each VF within each sample type and sorted by the frequency in corals.

Coral and seawater phages also encoded 71 unique genes classified as virulence factors (VFs) based on their role in virulence in known pathogens (62). It is important to note that while these genes are involved in virulence in known pathogens, they may have other roles in the molecular bacteria-eukaryotic interactions in commensal or mutualistic relationships (62). 51 of these VFs were unique to coral phages, 13 were shared between corals and seawater, and only 7 were unique to the seawater phages (Fig. 3C). The majority (70.00%) of VFs in coral phages encoded proteins related to effector delivery systems. This was followed by VFs involved in stress survival (10.29%) and immune modulation (9.14%). Seawater phages primarily encoded VFs involved in adherence (35.92%) and immune modulation (19.42%), but also encoded VFs related to stress survival at a similar frequency as in coral viruses (11.65%). The most abundant VF gene in corals was *ankY/legA9,* encoding for a type IV secretion system (T4SS) effector and accounting for 22.86% of VFs in coral phages (Fig. 3D). Coral phages carried a variety of other VF genes, such as *pipB2* (T2SS effector; 11.42%) and *icmE* (DotG T4SS central channel protein; 11.14%), which are also components of effector delivery systems. *csgG*, involved in curli production, was the most abundant VF gene in seawater phages (32.04%).

### Coral phages preferentially infect Alphaproteobacteria, Bacteroidia, and Halanaerobiia

To link viruses with their hosts, we binned and taxonomically classified 316 bacterial MAGs (bMAGs) from this metagenomic dataset (belonging to 41 bacterial classes) and combined them with the 310 publicly available bacterial isolate genomes from coral (5 classes) for a total of 626 putative bacterial hosts. We identified 299 putative links between 144 hosts and 275 GCVDB viruses based on the presence of proviruses (N = 236) and CRISPR spacer matches (N = 63) (Table S6). In total, 24.2% of bacterial isolates (N = 75) and 15.5% of bMAGS (N = 49) contained at least one provirus. Alphaproteobacteria, the most abundant bMAG class (28.80%), was overrepresented among phage-linked bMAGs (53.33% of CRISPR spacer links and 54.24% of provirus links; Fig. 4A). Gammaproteobacteria were the second most abundant class of bMAGs representing 15.19% and were underrepresented in their CRISPR spacer (6.67%) and provirus (8.47%) links. The classes Halanaerobiia and Bacteroidia, both comprising of obligate anaerobes, represented less than 13% of the bMAGS combined, yet had over twice as many CRISPR spacer linkages as Gammaproteobacteria bMAGs. Among bacterial isolates, however, Alphaproteobacteria links were underrepresented, accounting for only 5.56% of CRISPR spacer links and 41.24% of provirus links, and Gammaproteobacteria isolates were overrepresented in their linkages, accounting for 94.44% of CRISPR spacer linkages and 55.37% of provirus linkages (Fig. 4B). We also identified, for the first time to the best of our knowledge, viruses predicted to infect the putative coral mutualist *Endozoicomonas,* which harbored more than one prophage per genome (Tables S6).

**Figure 4.**
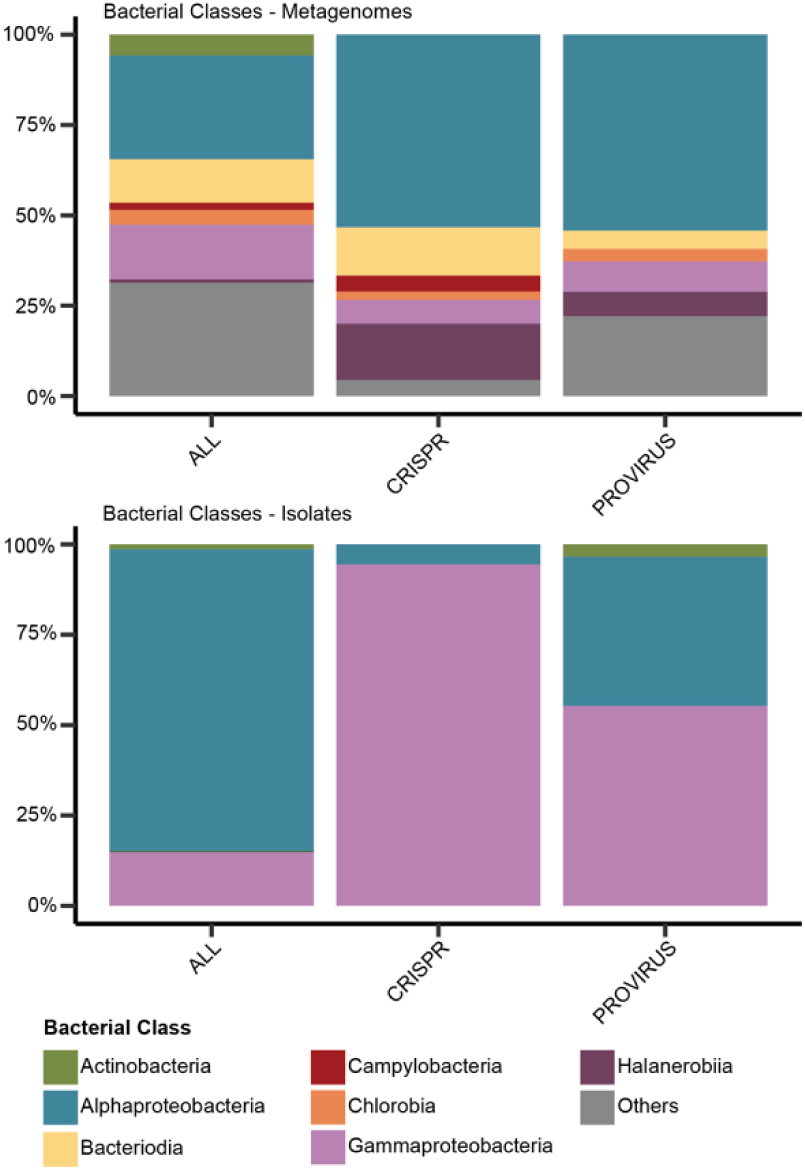
Bacteriophage-bacteria interactions. Frequency of **(A)** bMAGS and **(B)** isolates in the metagenomic dataset (ALL) and within those with CRISPR spacer or prophage linkages, at the taxonomic level of class. Classes with abundance < 2% (except for *Halanaerobiia*, which represented < 1% of bMAGs, but 7-16% of linkages) were grouped as “Others”.

### Tripartite network

Links between viruses and their putative hosts (N = 299) and the genes encoded by viruses in the GCVDB (N = 5,219) were used to construct a bacteria-phage-gene network (Fig. 5A). Among bacteria, the average clustering coefficient, which describes the connections between neighbors of a node, was 0.00., i.e., viruses linked to a bacterial host were not linked to each other based on shared metabolic and virulence genes. The top six ranks of predicted keystone viruses included 17 viruses (Table S10), all of which were classified as phages in the realm *Duplodnaviria* without family names or host linkages. These viruses primarily encoded the most common genes within the GCVDB, such as *DNMT1, dcm*, *DNMT3A*, and *ankY/legA9*. Importantly, the highest-ranking viruses encoded both metabolic and virulence genes, which reflected on their closeness centrality scores within the network.

**Figure 5.**
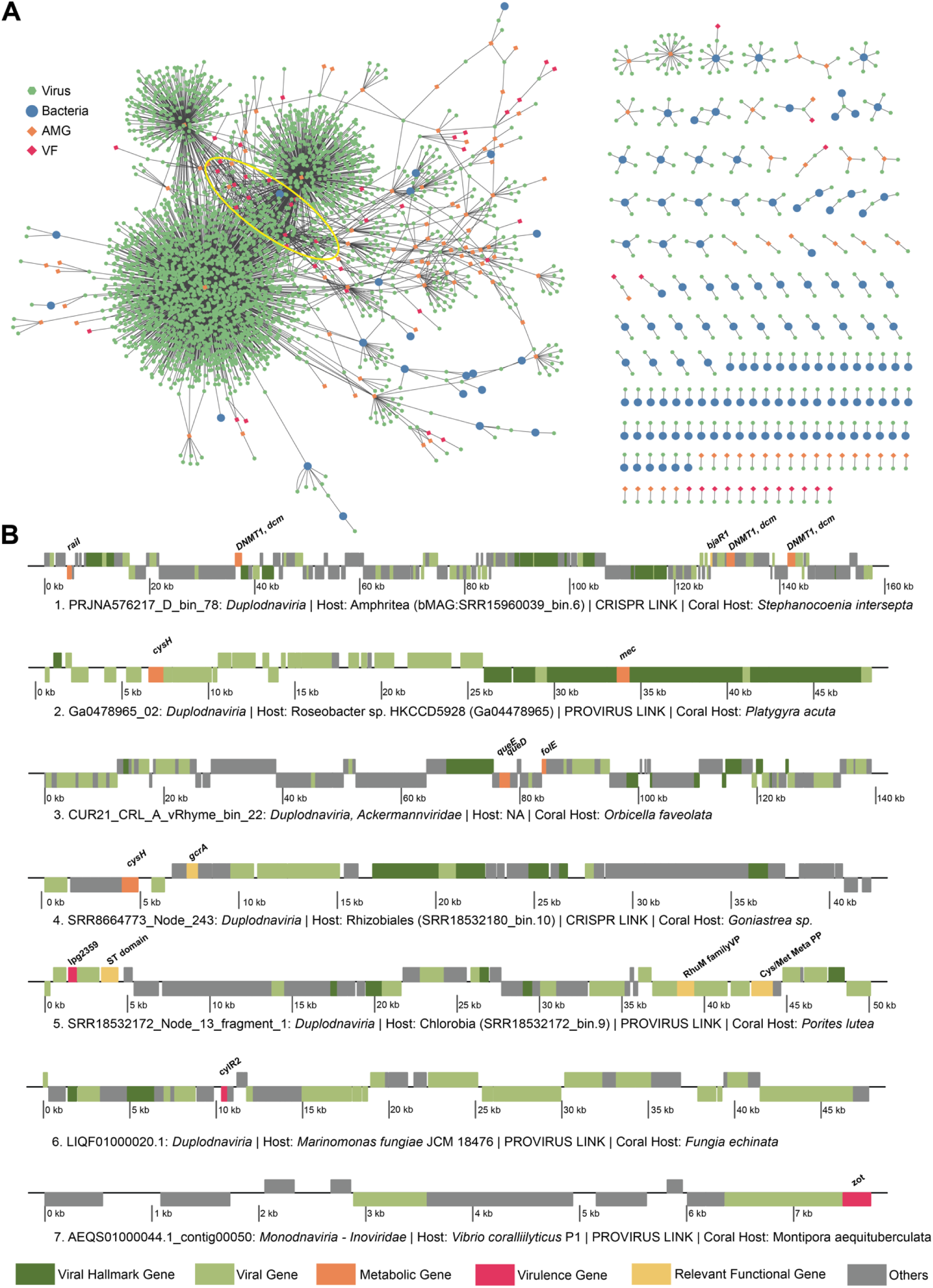
Bacteria-phage-gene network. **(A)** GCVDB viruses were connected to bacterial hosts (isolate and bMAG) according to the host prediction described in the methods and to metabolic genes and virulence factors according to their gene annotations. The yellow ellipse indicates the position of viruses identified as keystones in the network analyses. **(B)** Genome annotations of seven viral genomes of interest due to their host link or the presence of metabolic or virulence genes. The colors indicate gene classifications.

Among viruses with within the network, we show seven which encode ecologically relevant metabolic or virulence genes, six of which have been linked to bacterial hosts (Fig. 5B). Among the *Duplodnaviria* viruses, PRJNA576217_D_bin_78 (1) is CRISPR spacer linked to bMAG SRR15960039_bin.6 belonging to the genus *Amphritea,* family *Oceanospirillaceae* of *Gammaproteobacteria*. This virus encoded DNA cytosine-5 methyltransferase 1 (*DNMT1, dcm*), acyl-homoserine-lactone synthase (*raiI)*, and quorum-sensing system regulator (*BjaR1*). Another *Duplodnaviria*, Ga0478965_02 (2), was integrated into the genome of *Roseobacter* sp. HKCCD5928 (Alphaproteobacteria) isolated from the coral *Platygyra acuta*. This virus-encoded phosphoadenosine phosphosulfate reductase (*cysH)* and CysO sulfur-carrier protein-S-L-cysteine hydrolase (*mec*). CUR21_CRL_A_vRhyme_bin_22 (3), identified as a *Duplodnaviria* virus in the family *Ackermannviridae*, was not linked to a host, but encoded Queuosine biosynthesis genes which are of known importance to the phage-host evolutionary arms race. queD, a 6-pyruvoyltetrahydropterin/6-carboxytetrahydropterin synthase, queE, a 7-carboxy-7-deazaguanine synthase, and GCH1, folE, a GTP cyclohydrolase all play roles in deazaguanine DNA modifications in phages. SRR8664773_Node_243 (4) was CRISPR spacer linked to a *Rhizobiales* bMAG (SRR18532180_bin.10). SRR8664773_Node_243, from a *Goniastrea minuta* metagenome, encoded metabolic gene *cysH*, as well as bacterial cell cycle regulator GcrA. SRR18532172_Node_13_fragment_1 (5) a *Duplodnaviria* virus derived from *Porites lutea* skeleton, was integrated in a bMAG (SRR18532172_bin.9) from the same sample. This virus encoded a Sulfotransferase domain (ST domain), Cysteine and Methionine metabolism PLP-dependent enzyme (Cys/Met Meta PP) as well as virulence factors, including a type IV secretion system effector (lpg2359) and a RhuM family virulence protein (RhuM family VP). *Duplodnaviria* virus LIQF01000020.1 (6) was identified in the genome of *Marinomonas fungiae* JCM18476 isolated from the coral host *Fungia enchiata* and encoded the virulence factor *cylR2*, a cytolysin regulator. The isolated coral pathogen *Vibrio coralliilyticus* strain P1 carried an integrated *Monodnaviria* virus in the *Inoviridae* family of chronic filamentous viruses (AEQS01000044.1_contig00050; 7) encoding the gene for the zonula occludens toxin (*zot*).

## Discussion

Here, we introduce a size fractionation method to significantly increase the recovery of viral and bacterial DNA in coral metagenomes. By combining data generated across worldwide coral metagenomic and culture-based studies, including 710 coral metagenomes, viromes, and bacterial isolates, we identified 531 high- and medium-quality metagenome-assembled viral genomes from corals. These viruses infect diverse hosts and encode genes that contribute to holobiont functioning, specifically in Sulfur-containing amino acid metabolism and effector production and delivery.

### Enrichment of Viruses and Bacteria in coral metagenomes

VBE captured high and medium-quality viral genomes across four of the six viral realms recognized by the ICTV, with the exception of Adnaviria and Ribozyviria (Fig 2B). By using size-fractionation and sequencing viruses and bacteria together, the VBE method incurred fewer compositional biases compared to previous methods (63). Chloroform treatment efficiently removes bacteria, but selects against enveloped viruses, some non-enveloped viruses, and a third of tailed phages (63–65). Cesium chloride (CsCl) gradients enhance viral recovery (63) but severely bias viral diversity due to differences in viral capsid density (66) and often require the use of amplification methods that incur further biases (67). In viral studies that avoided these biases, viral genomes were not be assembled, limiting the analyses to single reads and compromising diversity and functional interpretations (10,25). Here, bacterial and viral genome recovery through VBE was possible even at a relatively low sequencing coverage (37 million reads and 5.5 billion bases per sample on average). This recovery represents an improvement from previous studies that required coverages from 58 to 146 million reads (68–71) to assemble bacterial MAGs and were most successful with samples devoid of coral tissue, such as in studies focused on the skeleton communities (68). VBE uses a coral fragment not much larger than a parrotfish bite, which is minimally invasive to corals. Therefore, this method could be easily adapted for the recovery of bacterial and viral RNA, which would presumably require larger amounts of input material (72). This method also pools all holobiont compartments (skeleton, tissue, and mucus), and adaptations are required for application to specific compartments (69,73). Increasing the size of the coral input sample and sequencing depth, and incorporating long-read data, may lead to the resolution of more complete genomes covering a larger biological diversity.

### An updated taxonomy of coral-associated viruses

The discontinuation of morphology-based viral taxonomic assignments by the International Committee on Taxonomy of Viruses (ICTV) in 2022 has implications for studying bacterial viruses in corals. This taxonomic reorganization included the creation or relocation of one order, 22 families, 30 subfamilies, 321 genera, and 862 species while simultaneously eliminating families such as Podoviridae, Siphoviridae, and Myoviridae, along with the order Caudovirales (43). In previous studies, these families were consistently the most abundant members of the coral holobiont (9,10,20). Within the GCVDB, we identified viruses in the realms *Duplodnaviria*, *Monodnaviria*, *Varidnaviria*, and *Riboviria*. Among the phages, the vast majority belonged to the class *Caudoviricetes*, which includes all tailed bacterial and archaeal viruses with icosahedral capsids and a dsDNA genome (43). The most abundant families under new taxonomy were *Kyanoviridae*, and *Autographiviridae* which represented 65% of *Caudoviricetes*. The family *Kyanoviridae* likely represents many of the since-abolished Myoviridae phages previously identified in corals. These “T4-like” phages share the myovirus morphotype and exclusively infect Cyanobacteria, but did not have host linkages in our dataset (43). *Autographiviridae* viruses, which typically exhibit a lytic lifestyle, are characterized by podophage morphology, and are known to primarily infect Gammaproteobacteria hosts (74).

### Coral viruses encode a limited but distinct repertoire of metabolic genes

Through incorporation and expression of metabolic genes, phages can impact the function and ecological relationships of their hosts (75,76). Despite a massively higher sampling effort of corals (N = 400) in comparison with seawater (N = 18), coral phages had a comparatively lower frequency and a less diverse repertoire of metabolic genes, primarily encoding genes involved in amino acid metabolism. These results suggest strong selection of a few metabolic genes in phage genomes associated with corals. DNA (cytosine-5)-methyltransferase 1 (*DNMT1, dcm*) and DNA (cytosine-5)-methyltransferase 3A (*DNMT3A*), comprising 60.35% of the metabolic genes in coral viruses, are involved in DNA methylation for evasion of restriction-mediated resistance against phage infection (77). They were found in viruses infecting eight different bacterial families, indicating widespread evolutionary arms race between coral viruses and their bacterial hosts (78). Another possible function of DNMT1, dcm is in the colonization of mucosal surfaces. In Group B *Streptococcus* specifically, knockout of *dcm* was found to reduce binding to immobilized mucin (79). We predict that in corals, these genes are involved in evasion of bacterial defense during lytic infections and possibly in enhancing bacterial host mucus colonization. S – (hydroxymethyl) glutathione dehydrogenase (*frmA, ADH5, adhC*), the next most frequent metabolic gene in coral viruses, plays a role in formaldehyde metabolism, which could be important for biomass and energy production of methylotrophs and other fermenting bacteria (80). Corals display significant diurnal oxygen fluctuations, with daytime photosynthetic oxygen production followed by rapid consumption by heterotrophic metabolism at night and resultant shifts to anaerobic metabolism (81). Viruses carrying *frmA, ADH5, adhC* genes, if infecting anaerobes which proliferate during nocturnal respiration, may bolster downstream production of formate, a key electron donor in anaerobic respiration (82). The contribution of fermentation to coral holobiont metabolism during nighttime oxygen depletion and the phage contribution to this shift should be the focus of future studies.

### Bacteria-eukaryote interaction genes are rare in coral phage genomes

Previous studies indicated that approximately 40% of viral sequences from corals were related to bacterial pathogenicity, annotated as “Virulence, Disease, and Defense” (10). While these genes are involved in pathogenicity on known pathogens, they can be involved in commensal or mutualistic bacteria-eukaryote interactions in non-pathogens (62). Here, 3.3% of phage genomes in the GCVDB and 2.2% of phage genomes in the seawater samples encoded at least one virulence factor. The frequency of virulence genes found in GCVDB and the seawater viruses in this study more closely aligned with the frequency observed in coral-seawater boundary layer viruses (58), where 2-4% of the viral community encoded virulence genes. This order of magnitude difference may be due to the use of a read-based bioinformatic approach in the study that found higher frequencies (10) in contrast to an assembly and MAG binning approach used here. The functions of these virulence factors also differed between studies. Here, we observed a broader diversity of VFs in coral phages relative to those in seawater phages. Seawater phages often encoded genes related to adherence and invasion, consistent with the coral boundary layer samples (58). In contrast, most virulence factors encoded by coral viruses were associated with effector delivery systems, whether encoding for effectors themselves (ankY/legA9; Icm type IV secretion system effector) or parts of the systems (icmE/dotG; Icm type IV secretion system central channel protein). Effector delivery systems enable the direct translocation of effectors into host cells through an injection apparatus (83). The predominant VFs in coral viruses, type IV secretion systems (T4SSs), play crucial roles in the virulence of pathogenic bacteria by facilitating interactions with prokaryotic competitors and eukaryotic hosts by translocating toxic effector proteins directly into other cells (83,84). Greater microbe-microbe competition in the dense coral microbiome (85) and frequent bacteria-animal interactions may drive the selection of these secretion systems in the phages identified here.

### Host links and network

Associating uncultivated viruses with their microbial hosts remains a challenge not only for coral microbiomes but also for any phage-host studies (49,57). Here, our ability to identify bacterial MAGs enabled, for the first time to the best of our knowledge, to predict virus-host pairs through CRISPR matches and integrated proviruses. The most abundant bMAG hosts, Alphaproteobacteria and Gammaproteobacteria, displayed an interesting trend where Alphaproteobacteria hosts were overrepresented among bMAGs with linkages and Gammaproteobacteria hosts were underrepresented, suggesting higher frequency of viral infection in the more abundant Alphaproteobacteria, consistent with observations of Alphaproteobacteria in coral reef seawater (57). Lower completeness of Gammaproteobacterial genomes (∼7% less complete than Alphaproteobacteria bMAGs, t-test, t(137) = 2.51, P = 1.34e-02) partially, but not fully explains the underrepresentation of Gammaproteobacteria linkages. This may simply indicate lower frequency of viral infections of Gammaproteobacteria in corals for unknown reasons. Halanaerobiia and Bacteroidiia constituted a minor fraction of bMAGs, yet they represented a substantial percent of the phage linkages. This observation aligns with a previous investigation in coastal seawater environments, wherein an abundance of free rRNA, indicative of recently lysed cells, was noted for copiotrophic and low-abundance bacteria (86). bMAGs in these groups had multiple phage linkages, suggesting that these taxa are susceptible to infection by multiple viruses.

The network analysis revealed several keystone *Duplodnaviria* viruses. Keystone viruses shared a diversity of metabolic genes (which makes them both more connected in the network and more able to diversify the functional capacity of bacteria within the holobiont). Low clustering across bacterial members of the network shows that the viruses infecting the same bacterial host do not share metabolic genes. This indicates that different viruses create distinct virocell metabolism upon infection of the same host (87). A closer inspection of viral genomes with connections with hosts and genes of interest revealed the genomic underpinnings of phage-bacteria-coral interactions. *Duplodnaviria* phage PRJNA576217_D_bin_78 encodes an acyl-homoserine-lactone (AHL) synthase (*raiI*) and a LuxR family transcriptional regulator (*bjaR1)*, which are both involved in quorum sensing. RaiI produces AHL, which upon reaching a critical threshold within a bacterial cell, triggers AHL-signaling and the initiation of the expression of virulence genes (88). BjaR1 acts as a quorum-sensing transcriptional regulator involved in the response to AHL (89). The presence of both genes in the same phage suggests this phage coopts the AHL signaling of its bacterial host (bMAG SRR15960039_Bin.6) to modulate the expression of virulence genes and/or direct lysis-lysogeny decisions. The host, *Amphritea sp*, is an Oceanospirillaceae in the Gammaproteobacteria class (90). A bMAG of the same genus was found across metagenomes of a stony coral tissue loss diseased *Stephanocoenia intersepta* in the original study (69) and has been found in association with black band disease (91). We speculate that the virus identified here is involved in the pathogenicity of *Amphritea* in corals through modulation of quorum sensing signaling (Fig. 6A).

**Figure 6.**
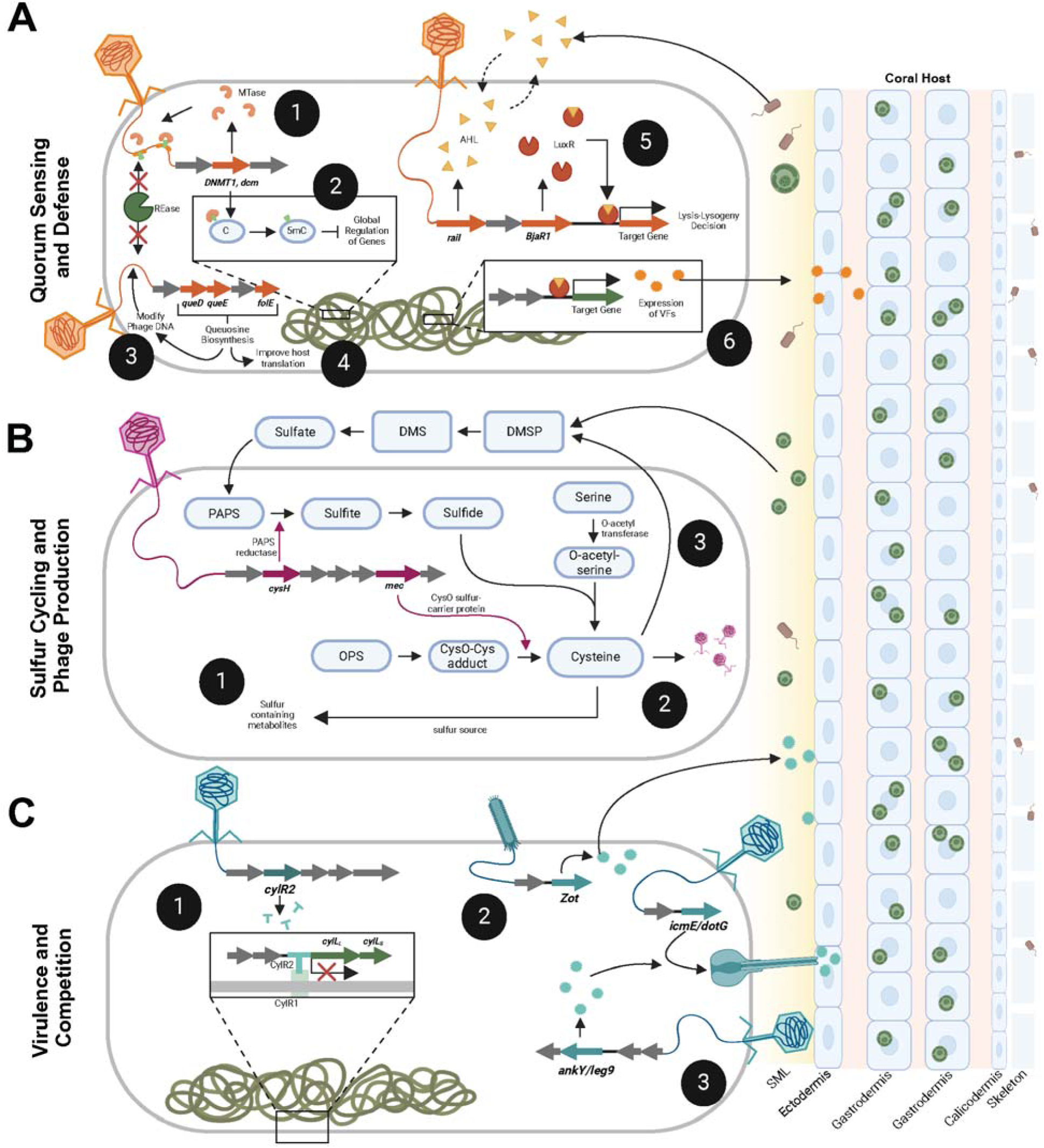
Conceptual hypotheses for mechanisms underpinning phage roles on coral-associated microbiomes. **(A)** Viral infection and gene transfer impact bacterial quorum sensing and defense where phage-encoded methyltransferases (1) protect phage nucleic acids from host-encoded restriction endonucleases and (2) regulate adhesins and metabolic factors with putative roles in carbohydrate metabolism and mucin adherence. Genes involved in the synthesis of Queuosine precursors may be used to (3) modify phage DNA as protection against host restriction systems or (4) increase Queuosine levels in host tRNAs, improving the host’s translational efficiency. Quorum-sensing genes encoded by phages may influence (5) lysis-lysogeny decisions and (6) the expression of genes related to eukaryotic host interactions (virulence genes). **(B)** Phages may contribute to Sulfur cycling through (1) contributing to the production of sulfur-containing metabolites that are effective against oxidative stress, (2) the diversion of Sulfur towards Cysteine production and assimilation into phage proteins, and (3) interference in the synthesis of DMSP due to the deviation of the DMSP precursor cysteine towards phage proteins. **(C)** Phages may impact virulence and competition within the coral holobiont by (1) regulating the expression of bacterial cytolysins, (2) expression of exotoxins that increase the permeability of coral host tissues, and (3) encoding type IV secretions system central channel proteins and effectors that can mediate bacteria-bacteria or bacteria-eukaryote interactions.

Evidence of the phage-host evolutionary arms race was present in several genomes, as many viruses encoded DNMT1, dcm, likely for evasion of restriction-mediated resistance. Another mechanism of this restriction-evasion was demonstrated in the genome CUR21_CRL_A_vRhyme_bin_22 (3) The Queuosine biosynthesis genes encoded by this phage, queD, queE, and folE, are widespread in phages, especially in the genomes of those with pathogenic hosts (92). These genes are suggested to modify phage DNA as protection against host restriction systems and contribute to the level of Queuosine in host tRNAs, improving host translational efficiency (92).

Roseobacters constitute a significant portion of the coral mucus microbiome, and while the precise nature of their interactions with coral remains ambiguous, there is evidence suggesting their potential roles as probiotics in coral reproduction and defense (93). The degradation and production of DMSP/DMS by roseobacters is considered a nutrient source for corals (94,95). Here, a phage genome (Ga0478965_02) within a Roseobacter isolate (sp. HKCCD5928) encoded metabolic genes related to Sulfur metabolism, specifically assimilatory sulfate reduction (phosphoadenosine phosphosulfate reductase; *cysH*) and the transfer of sulfur during cysteine production (CysO sulfur-carrier protein; *mec*). Viruses commonly utilize sulfur metabolism genes to increase the rates of cysteine production for viral particle assembly (42). Yet, these sulfur genes were encoded in a prophage, Ga0478965_02. It is possible that these genes are transcribed only when the prophage enters the lytic cycle (96). Alternatively, its expression during lysogeny contributes to the synthesis of DMSP and sulfur-containing metabolites by the Roseobacter host (Fig. 6B). Other bacterial hosts associated with the skeletal components of coral hosts exhibited connections to viruses that carried multiple genes associated with the metabolism of sulfur-containing amino acids and metabolites, exemplified by instances like SRR8664773_Node_243 and SRR18532172_Node_13_fragment_1. While these phage-encoded genes could potentially play a role in dimethylsulfoniopropionate (DMSP) cycling and phage production, the production of sulfur-containing metabolites has been previously linked to resistance against oxidative and nitrosative stress in various bacterial hosts (97).

Genes related to effector delivery, with potential roles in bacterial competition and interactions with the coral host, were the most common “virulence factors” in the GCVDB. Often, these phage-encoded effector toxin proteins are multifunctional, but they can convert bacterial hosts from non-pathogenic to virulent (98) (Fig. 6C). The presence of these genes in our dataset indicates a potentially important role of phages in contributing to bacterial virulence in corals, perhaps contributing to coral disease. One of the viral genomes identified here, *Duplodnaviria* virus LIQF01000020.1, encodes genes involved in the regulation of cytolysin, an exotoxin that lyses prokaryotic and eukaryotic cells (99). A prophage encoding zonula occludens toxin (Zot) previously detected in the coral pathogen *Vibrio coralliilyticus* strain P1 was also observed here. These observations support the idea that certain coral diseases may result from lysogenic conversion (100,101). However, these “virulence” genes can also be involved in other non-pathogenic functions in the molecular interactions between bacteria and eukaryotic cells when they are called “fitness factors” (62). Here, these cells may include the coral animal cells, the endosymbiotic algae, or other eukaryotes in the holobiont.

## Conclusion

The results described here reveal the basis of mechanisms by which bacteriophages interact with bacteria within the coral holobiont, with potential downstream effects on bacteria-bacteria and bacteria-coral interactions. The large dataset and high resolution gained from the assembly and binning of genomes enabled the use of stringent quality thresholds, increasing confidence in the annotations and predictions made. The gene-resolved phage-host network highlighted phages’ selective repertoire of metabolic genes that can impact bacterial communication, competition, and molecular interactions with the coral host, illuminating the genomic underpinnings of phage-bacteria-coral symbioses.

## Data availability

Coral metagenomic sequence data generated for this project are available on NCBI (PRJNA1061506 and PRJNA975592). The code used to conduct this analysis is publicly available on GitHub (https://github.com/Silveira-Lab/Wallace_Coral_Holobiont_Viruses). Assembled viral genomes are available on Figshare: https://doi.org/10.6084/m9.figshare.24968805.v1. Viral genomes with multiple contigs are N-linked. Bacterial MAGs are available on Figshare: https://doi.org/10.6084/m9.figshare.25000160.v1.

## Authors’ contributions

CS conceptualized and led the development of the VBE method. CS and BW conceptualized the study. BW and NV conducted sampling, method development, and sample processing. BW conducted data retrieval, pipeline development, and analyses. PHB conducted analyses. BW wrote the first draft, and all authors edited the manuscript.

